# A Chromosome-level Sequence Assembly Reveals the Structure of the *Arabidopsis thaliana* Nd-1 Genome and its Gene Set

**DOI:** 10.1101/407627

**Authors:** Boas Pucker, Daniela Holtgräwe, Kai Bernd Stadermann, Katharina Frey, Bruno Huettel, Richard Reinhardt, Bernd Weisshaar

## Abstract

**Background:** In addition to the BAC-based reference sequence of the accession Columbia-0 from the year 2000, several short read assemblies of THE plant model organism *Arabidopsis thaliana* were published during the last years. Also, a SMRT-based assembly of Landsberg *erecta* has been generated that identified translocation and inversion polymorphisms between two genotypes of the species.

**Results:** Here we provide a chromosome-arm level assembly of the *A. thaliana* accession Niederzenz-1 (AthNd-1_v2c) based on SMRT sequencing data. The best assembly comprises 69 nucleome sequences and displays a contig length of up to 16 Mbp. Compared to an earlier Illumina short read-based NGS assembly (AthNd-1_v1), a 75 fold increase in contiguity was observed for AthNd-1_v2c. To assign contig locations independent from the Col-0 gold standard reference sequence, we used genetic anchoring to generate a *de novo* assembly. In addition, we assembled the chondrome and plastome sequences.

**Conclusions:** Detailed analyses of AthNd-1_v2c allowed reliable identification of large genomic rearrangements between *A. thaliana* accessions contributing to differences in the gene sets that distinguish the genotypes. One of the differences detected identified a gene that is lacking from the Col-0 gold standard sequence. This *de novo* assembly extends the known proportion of the *A. thaliana* pan-genome.

## Background

### Introduction

*Arabidopsis thaliana* became the most important model for plant biology within decades due to properties valuable for basic research like short generation time, small footprint, and a small genome [1]. Shortcomings of the BAC-by-BAC assembled 120 Mbp long Col-0 gold standard sequence [2] are some missing sequences and gaps in almost inaccessible regions like repeats in the centromeres [3, 4], at the telomeres and throughout NORs as well as few mis-assemblies [5, 6]. Information about genomic differences between *A. thaliana* accessions were mostly derived from short read data [7-9]. Only selected accessions were sequenced deep enough and with sufficient read length to reach almost reference-size assemblies [7, 10-15]. While the identification of SNPs can be based on short read mappings, the identification of structural variants had an upper limit of 40 bp for most of the investigated accessions [9]. Larger insertions and deletions, which will often result in presence/absence variations of entire genes, are often missed in short read data sets but are easily recovered by long read sequencing [14-16]. *De novo* assemblies based on long sequencing reads are currently emphasized to resolve structural variants without an upper limit and to facilitate *A. thaliana* pan-genomics. Even a fully complete Col-0 genome sequence would not reveal the entire diversity of this species, as this accession is assumed to have a relatively small genome compared to other *A. thaliana* accessions.

The strong increase in the length of sequencing reads that was technically realized during the last years is enabling new assembly approaches [17, 18]. Despite the high error rate of ‘Single Molecule, Real Time’ (SMRT) sequencing, the long reads significantly improve contiguity of *de novo* assemblies due to an efficient correction of the almost unbiased errors [19-21], provided that sufficient read coverage is available. SMRT sequencing offered by PacBio results routinely in average read lengths above 10 kbp [10, 22, 23]. These long reads were incorporated into high quality hybrid assemblies involving Illumina short read data [14, 23], but increasing sequencing output supports the potential for so called ‘PacBio only assemblies’ [10-12, 15, 20]. Oxford Nanopore Technology’s (ONT) sequencing provides even longer reads with recent reports of longest reads over 2Mbp [24]. However, the error rate of 5-15% [25] is still an issue in plant genome assembly although it has been shown that a high contiguity assembly is possible for *A. thaliana* [15].

Here we provide a SMRT sequencing-based *de novo* genome assembly of Nd-1 comprising contigs of chromosome-arm size anchored to chromosomes and oriented within pseudochromosome sequences based on genetic linkage information. The application of long sequencing reads abolished limitations of short read mapping and short read assemblies for genome sequence comparison. Based on this genome sequence assembly, we identified genomic rearrangements between Col-0 and Nd-1 ranging from a few kbp up to one Mbp. Gene duplications between both accessions as well as lineage specific genes in Nd-1 and Col-0 were revealed by this high quality sequence. The current assembly version outperforms the Illumina-based version (AthNd-1_v1) about 75 fold with respect to assembly contiguity calculated with respect to number of contigs [13] and is in the same quality range as the recently released L*er* and KBS-Mac-74 genome sequence assembly [14, 15].

## Methods

### Plant material

Niederzenz-1 (Nd-1) seeds were obtained from the European Arabidopsis Stock Centre (NASC; stock number N22619). The DNA source was the same as described earlier [13].

### DNA extraction

The DNA isolation procedure was based on previously published protocols [12, 26] and started with 5 g of frozen leaves which were homogenized by grinding. Samples were mixed in a 1:10 ratio with extraction buffer (300mM Tris pH8.0, 25mM EDTA, 2M NaCl, 2% polyvinylpyrrolidone (PVP), 2% hexadecyltrimethylammonium bromide (CTAB)) and incubated at 65°C for 30 minutes with six inversions to mix the samples again. After five minute spinning at 5,000xg, the supernatant was transferred and mixed with one volume of chlorophorm/isoamylalcohol (24:1). Again, the upper phase was transferred after repetition of the centrifugation step for ten minutes. RNA was removed by adding 30μL RNase A (10mg/ml) and incubation for 30 minutes at 37°C. Addition of chlorophorm/isoamylalcohol, centrifugation and transfer of supernatant were repeated. One volume of isopropanol and 0.1 volumes of 3M NaOAc (pH 5.2) were added and mixed. DNA was precipitated by incubating at −80°C for 30 minutes and spinning for 45 minutes at 5,000g and a final ethanol wash step was performed. Finally, 500μl of 10mM Tris/HCl (pH 8.0) were added and samples were incubated over night at 4°C for resuspension.

### Library preparation and sequencing

Sequencing was performed using PacBio RS II (Menlo Park, CA, USA). Five microgram high molecular weight DNA without further fragmentation was used to prepare a SMRTbell library with PacBio SMRTbell Template Prep Kit 1 (Pacific Biosciences, Menlo Park, CA, USA) according to the manufacturer’s recommendations. The resulting library was size-selected using a BluePippin system (Sage Science, Inc. Beverly, MA, USA) to enrich for molecules larger than 11 kbp. The recovered library was again damage repaired and then sequenced on a total of 25 SMRT cells with P6-C4v2 chemistry and by MagBead loading on the PacBio RSII system (Pacific Biosciences, Menlo Park, CA, USA) with 360 min movie length.

### Assembly parameters

A total of 1,972,766 subreads with an N50 read length of 15,244 bp and containing information about 16,798,450,532 bases were generated. Assuming a genome size of 150 Mbp, the data cover the genome at 112 fold.

Read sequences derived from the plastome [GenBank: AP000423.1] or chondrome [GenBank: Y08501.2] were extracted from the raw data set by mapping to the respective sequence of Col-0 as previously described [27]. Canu v1.4 [28] was used for the assembly of the organell genome sequences. Beside default parameters, a genome size of 0.37 Mbp for the chondrome and 0.167 Mbp for the plastome were assumed in the assembly process. Scaffolding of initial contigs was performed with SSPACE-LongRead v1.1 [29] (parameters: -a 0 -i 70 -k 3 -r 0.3 -o 10). The quality of both assemblies was checked by mapping of NGS reads from Nd-1 [13] and Col-0 [30]. Manual inspection and polishing with Quiver v.2.3.0 (parameters: minSubReadLength=500, readScore>80, minLength=500, maxHits=10, maxDivergence=30, minAnchorSize=12, --maxHits=1, --minAccuracy=0.75, --minLength=50, --algorithmOptions=“-useQuality”, --seed=1, --minAccuracy=0.75, --minLength=50, --concordant --algorithmOptions=“-useQuality”, minConfidence=40, minCoverage=5, diploidMode=False) [12] let to the final sequences. The start of the Nd-1 plastome and chondrome sequences was set according to the corresponding Col-0 sequences to ease comparisons. Finally, small assembly errors were corrected via CLC basic variant detection (ploidy=2; coverage<100,000; minimalCoverage=5; minimalCount=5; minimalFrequency=0.2) based on mapped Illumina paired-end reads (SRX1683594, [13]) and PacBio reads. Sequence properties like GC content and GC skew were determined and visualized by CGView [31].

A total of 166,600 seed reads spanning 4,500,092,354 nt (N50 = 26,295 nt) and covering the expected 150 Mbp genome sequence were used for the assembly thus leading to a coverage of 30 fold (see AdditionalFile1 for details). Assemblies were performed with Canu [28], FALCON [12], Flye [32] and miniasm [33]. These assemblies are named Ath-Nd1_v2 with an assembler specific suffix (c=Canu, f=FALCON, y=Flye, m=miniasm).

All available SMRT sequencing reads were subjected to Canu v.1.7.1 [28] for read correction, trimming, overlap detection, and *de novo* assembly (see AdditionalFile2 for parameters). Contigs with a length below 50 kb were discarded to avoid artifacts. The remaining contigs were checked for contaminations with bacterial sequences and organell genome sequences as previously described [13]. BWA-MEM v0.7.13 [34] was used with default parameters and -m to map Nd-1 Illumina reads [13] to this assembly. The coverage depth was extracted from the resulting BAM file as previously described [35] and used to support the identification of plastome and chondrome contigs. Pilon v.1.22 [36] was applied twice with default parameters for polishing of the assembly.

Release version 1.7.5 of the FALCON assembler https://github.com/PacificBiosciences/FALCON/ [12] was used for a *de novo* assembly (see AdditionalFile3 for parameters) of the nuclear genome sequence. Resulting contigs were checked for contaminations with bacterial sequences and organell genome sequences as previously described [13]. Small fragments with low coverage were removed prior to polishing and error correction with Quiver [12].

SMRT sequencing reads were corrected via Canu and subjected to miniasm v0.3-r179 [33] for *de novo* assembly. Illumina reads were mapped with BWA-MEM for two rounds of polishing via Pilon as described above. The assembly was reduced to nucleome contigs as described above for the Canu assembly.

Flye v2.3.1 [32] was deployed on the subreads with an estimated genome size of 150 Mbp. The resulting assembly was polished twice via Pilon and reduced to nucleome contigs as described above for the Canu assembly.

### Construction of pseudochromosomes based on genetic information

All 26 contigs of Ath-Nd1_v2f were sorted and orientated based on genetic linkage information derived from 63 genetic markers (AdditionalFile4, AdditionalFile5, AdditionalFile6), which were analyzed in about 1,000 F2 plants, progeny of a reciprocal cross of Nd-1xCol-0 and Col-0xNd-1. Genetic markers belong to three different types: (1) fragment length polymorphisms, which can be distinguished by agarose gel electrophoresis, (2) small nucleotide polymorphisms which can be distinguished by Sanger sequencing and (3) small nucleotide polymorphisms, which were identified by high resolution melt analysis.

Design of oligonucleotides was performed manually and using Primer3Plus [37]. DNA for genotyping experiments was extracted from *A. thaliana* leaf tissue using a cetyltrimethylammonium bromide (CTAB) based method [38]. PCRs were carried out using GoTaq^®^ G2 DNA Polymerase (Promega) generally based on the suppliers’ protocol. The total reaction volume was reduced to 15 μl and only 0.2u of the polymerase were used per reaction. Sizes of amplicons generated were checked on agarose gels. If required, samples were purified for sequencing by ExoSAP-IT (78201.1.ML ThermoFisher Scientific) treatment as previously described [39]. Sanger sequencing on ABI3730XL was applied to identify allele-specific SNPs for the genotyping. Manual inspection of gel pictures and electropherograms lead to genotype calling. High resolution melt analysis was performed on a CFX96 Touch Real-Time PCR Detection System (BioRad) using the Precision Melt Supermix according to suppliers instructions (BioRad). All data were combined, processed by customized Python scripts to calculate recombination frequencies between genetic markers. Linkage of genetic markers provided information about relationships of assembled sequences. The north-south orientation of the chromosomes was transferred from the reference sequence based on RBH support. Contigs were joined to pseudochromosome sequences for Ath-Nd1_v2f. Subsequently, positioning was transferred to the 69 contigs of the Canu assembly to create pseudochromosome sequences for AthNd-1_v2c (AdditionalFile7).

Since the assemblies except Ath-Nd1_v2f contain smaller contigs in the range of 50-100 kb, anchoring via genetic linkage information was not feasible. Therefore, these short contigs were placed based on best BLASTn matches to the TAIR9 sequence (i.e. the Col-0 gold standard reference) and integrated into pseudochromosomes as previously described [13]. We used the TAIR9 sequence, because this is the sequence basis of the structural and functional annotation provided in TAIR9, TAIR10 and Araport11 [40]. Since the total length of the TAIR9 sequence exceeds that of the Ath-Nd1_v2f assembly, TAIR9 was selected for anchoring of small contigs.

All data generated is stored in the PGP Repository [41] and is accessible via DOI (https://doi.org/10.5447/IPK/2019/4).

### Genome structure investigation

Characteristic elements of the Nd-1 genome sequence were annotated by mapping of known sequences as previously described [13]. Fragments and one complete 45S rDNA unit were discovered based on gi|16131:848-4222 and gi|16506:88-1891. AF198222.1 was subjected to a BLASTn search for the identification of 5S rDNA sequences. Telomeric repeats were used to validate the assembly completeness at the pseudochromosome end as well as centromere positions as previously described [13].

### BUSCO analysis

BUSCO v3 [42] was run with default parameters on the Nd-1 pseudochromosomes and on the TAIR9 reference sequence to produce a gold standard for Arabidopsis. The ‘embryophyta_odb9’ was used as reference gene set.

### Genome sequence alignment

Nd-1 pseudochromosome sequences were aligned to the Col-0 gold standard sequence [2] via nucmer [43] and NucDiff [44] based on default parameters of NucDiff. Customized Python scripts were deployed to process the results and to assess the genome-wide distribution of differences. Spearman correlation coefficient was calculated using the implementation in the Python module scipy to validate the indication of increased numbers of SV around the centromeres.

### Gene prediction and RBH analysis

AUGUSTUS v.3.3 [45] was applied to all four Nd-1 assemblies with previously optimized parameters [39]. Afterwards, the identification of RBHs at the protein sequence level between Nd-1 and Col-0 (Araport11, representative peptide sequences) was carried out with a custom Python script as previously described [13]. Additionally, gene prediction was run on the Col-0 gold standard sequence [2] as well as on the L*er* chromosome sequences [14]. Parameters were set as described before to generate two control data sets. Previously trimmed and filtered ESTs [13, 46] were matched via BLASTn [47] to the predicted mRNAs. Pairwise global alignments were constructed via MAFFT v.7.299b [48] to validate the annotation quality. After processing GCA_000001735.1 (Col-0), Ath-Nd-1_v2c, and the most recent L*er* assembly [14] with RepeatMasker v4 [49] all three were subjected to cactus [50] for alignment. CAT [51] was run on this alignment with the ENSEMBL v42 annotation of Col-0.

INFERNAL [52] with default parameters and based on Rfam13 [53] was applied to detect various non-coding RNA genes. In addition, tRNAscan-SE [54] with –G option was deployed to identify tRNA and rRNA genes. Overlaps between both methods were analysed.

### Transposable element annotation

All annotated transposable element (TE) sequences of Araport11 (derived from TAIR) [40] were mapped via BLASTn to the Nd-1 assembly AthNd-1_v2c and against the Col-0 gold standard sequence. The top BLAST score for each element in the mapping against the TAIR9 reference sequence was identified. All hits against Nd-1 with at least 90% of this top score were considered for further analysis. Overlapping hits were removed to annotate a final TE set. All predicted Nd-1 genes which overlapped TEs with more than 80% of their gene space were flagged as putative protein encoding TE genes. In addition, RepeatModeler v.1.0.11 [49] was deployed to identify novel TE sequences in the Ath-Nd-1_v2c assembly.

### Identification of gene space differences

Genes in insertions in Ath-Nd1_v2c were searched in the Col-0 gold standard sequence and vice versa. The BLAST results on DNA and peptide level indicated the absence of any truly novel genes thus indicating that presence/absence variants are the results of gene duplications. Apparent gene space differences could be caused by regions missing in the assembly. Therefore, Nd-1 (SRR2919279, SRR3340908, SRR3340909) [13] and Col-0 (SRR1810832, SRR1945757)[9, 30] Illumina sequencing reads were mapped to Ath-Nd1_v2c via BWA MEM [34] using the -m flag to discard spurious hits. The sequencing read coverage depth was calculated via bedtools by a customized Python script [35]. Average coverage per accession was calculated as the median of all coverage values. Per gene coverage was calculated as the median of all coverage values at positions in the respective gene and normalized to the average coverage of the respective accession. To correct for accession specific mapability differences caused e.g. by sequence divergence from Nd-1, the resulting values were additionally normalized to the median of all per gene ratios. Genes were considered as duplicated in Col-0 if their relative coverage in Nd-1 was below 50 % of the Col-0 value. Genes with Nd-1 values above 150 % of the Col-0 value were considered duplicated in Nd-1. These cutoff values were validated based on experimentally validated gene differences. Corresponding AGIs were identified via BLASTp to transfer the functional annotation of Araport11 if possible. Following this initial identification, putative TE genes were removed based on the annotation or the overlap with annotated TE sequences (AdditionalFile8), respectively.

Sequencing data of 1,135 *A. thaliana* accessions [9] were retrieved from the Sequence Read Archive, mapped to Ath-Nd1_v2c, and processed as described above. Accessions displaying an average coverage below 10x were excluded from the following identification of presence/absence variations. GeneSet_Nd-1_v2.0 genes were classified as dispensable if their relative coverage was below 0.1 in more than 100 accessions. Remaining genes were classified as core or TE genes depending on their overlap with TE features (see TE annotation).

### Validation of rearrangements and duplications

LongAmpTaq (NEB) was used for the generation of large genomic amplicons up to 18 kbp based on the suppliers’ protocol. Sanger sequencing was applied for additional confirmation of generated amplicons. The amplification of small fragments and the following procedures were carried out with standard polymerases as previously described [13].

### Investigation of collapsed region

The region around At4g22214 was amplified in five overlapping parts using the Q5 High Fidelity polymerase (NEB) with genomic DNA from Col-0. Amplicons were checked on agarose gels and finally cloned into pCR2.1 (Invitrogen) or pMiniT 2.0 (NEB), respectively, based on the suppliers’ recommendations. Cloned amplicons were sequenced on an ABI3730XL by primer walking. Sequencing reads were assembled using CLC GenomicsWorkbench (v. 9.5 CLC bio). In addition, 2×250 nt paired-end Illumina reads of Col-0 [30] were mapped to correct small variants in the assembled contigs and to close a small gap between cloned amplicons.

### Analysis of gaps in the Col-0 reference sequence

Flanking sequences of gaps in the Col-0 gold standard sequence were submitted to a BLASTn against the Ath-Nd-1_v2c genome sequence. Nd-1 sequences enclosed by hits of pairs of 30 kbp long flanking sequences from Col-0 were extracted. Homotetramer frequencies were calculated for all sequences and compared against the frequencies in randomly picked sequences. A Mann-Whitney U test was applied to analyze the difference between both groups.

## Results

### Nd-1 genome

Assemblies generated via Canu (Ath-Nd-1_v2c), FALCON (Ath-Nd-1_v2f), Flye (Ath-Nd-1_v2y) and miniasm (Ath-Nd-1_v2m) were compared based on numerous assembly statistics (Table 1). AthNd-1_v2f exceeds the previously reported assembly version AthNd-1_v1 by 2.5 Mbp, while reducing the number of contigs by a factor of about 200 to 26. AthNd-1_v2c is adding 6.9 Mbp to the previous assembly version (AthNd-1_v1), but at the expense of a higher number of contigs than AthNd-1_v2f. We selected AthNd-1_v2c that contains 69 contigs as the representative assembly which should be used for further comparison. Pseudochromosomes of AthNd-1_v2f were constructed truly *de novo* from 3-7 contigs based on genetic linkage information. All 26 contigs were anchored based on 63 genetic markers, but precise positioning around the centromeres of chromosome 4 and 5 was ambigious. Pseudochromosomes reach similar lengths as the corresponding chromosome sequences in the Col-0 gold standard sequence. Pseudochromosomes for AthNd-1_v2c were generated by transferring orientation and position of large contigs from AthNd-1_v2f. The Nd-1 genome sequence AthNd-1_v2c contains complete 45S rDNA units on pseudochromosome 3 as well as several fragments of additional 45S rDNA units on all other pseudochromosomes (Fig. 1). Centromeric and telomeric repeat sequences as well as 5S rDNA sequences were detected at centromere positions. Completeness of most assembled sequences was confirmed by the occurrence of telomeric repeat sequences (Fig. 1). The high assembly quality and completeness of AthNd-1_v2c is supported by the detection of 98.2% of all embryophyta BUSCO genes – even one more than detected in the Col-0 gold standard sequence (AdditionalFile9).

**Table 1:**
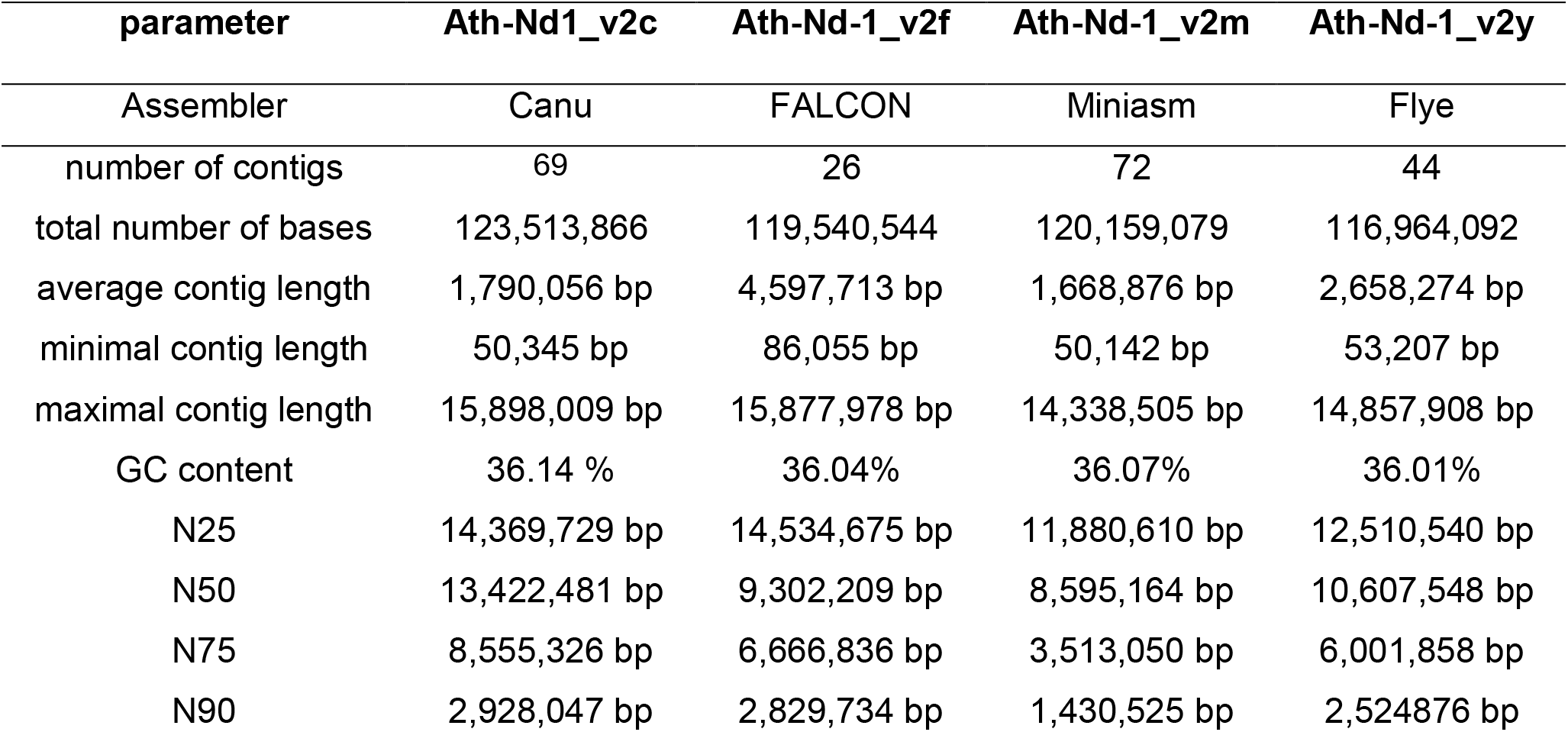
Nd-1 *de novo* assembly statistics. Metrics of assemblies of the Nd-1 nucleome sequence generated by Canu, FALCON, miniasm, and Flye. All described assemblies are the final version after polishing.

| parameter | Ath-Nd1_v2c | Ath-Nd-1_v2f | Ath-Nd-1_v2m | Ath-Nd-1_v2y |
| --- | --- | --- | --- | --- |
| Assembler | Canu | FALCON | Miniasm | Flye |
| number of contigs | 69 | 26 | 72 | 44 |
| total number of bases | 123,513,866 | 119,540,544 | 120,159,079 | 116,964,092 |
| average contig length | 1,790,056 bp | 4,597,713 bp | 1,668,876 bp | 2,658,274 bp |
| minimal contig length | 50,345 bp | 86,055 bp | 50,142 bp | 53,207 bp |
| maximal contig length | 15,898,009 bp | 15,877,978 bp | 14,338,505 bp | 14,857,908 bp |
| GC content | 36.14 % | 36.04% | 36.07% | 36.01% |
| N25 | 14,369,729 bp | 14,534,675 bp | 11,880,610 bp | 12,510,540 bp |
| N50 | 13,422,481 bp | 9,302,209 bp | 8,595,164 bp | 10,607,548 bp |
| N75 | 8,555,326 bp | 6,666,836 bp | 3,513,050 bp | 6,001,858 bp |
| N90 | 2,928,047 bp | 2,829,734 bp | 1,430,525 bp | 2,524,876 bp |

**Figure 1:**
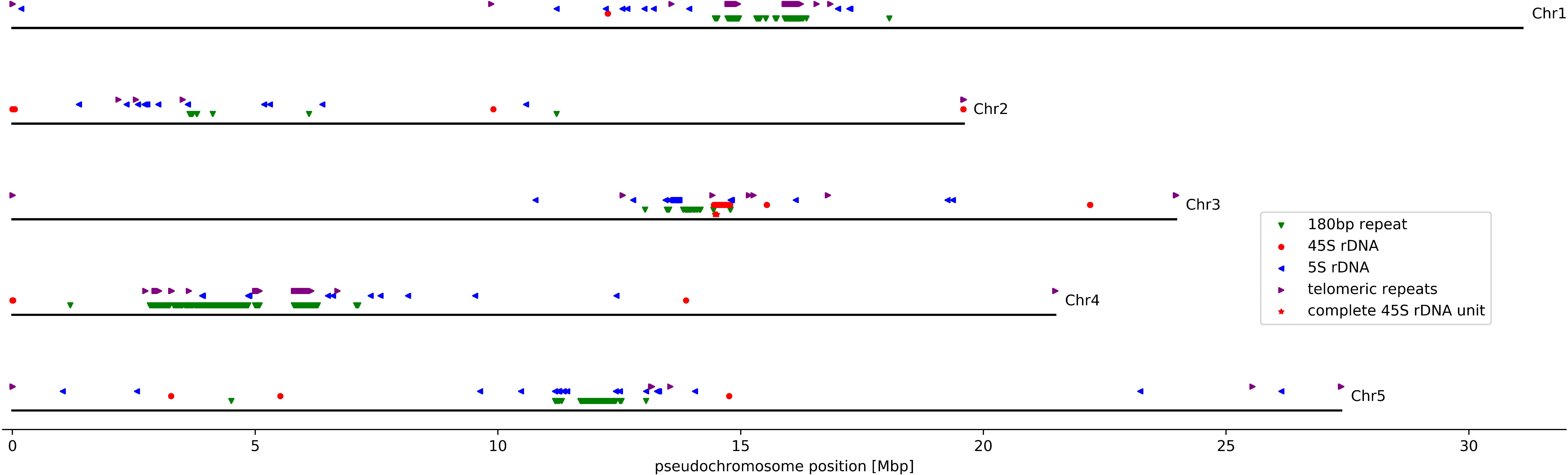
Nd-1 genome structure. Schematic pseudochromosomes are represented by black lines with positions of genomics features highlighted with colored icons as indicated in the insert.

The plastome and chondrome sequences comprise 154,443 bp and 368,216 bp, respectively (https://doi.org/10.5447/IPK/2019/4). A total of 148 small variants were identified from a global alignment between the Nd-1 and Col-0 plastome sequences. General sequence properties like GC content and GC skew (AdditionalFile10, AdditionalFile11) are almost identical to the plastome and chondrome of Col-0. Nevertheless, there are some rearrangements between the chondrome sequences of Nd-1 and Col-0.

### Genome structure differences

Sequence comparison between AthNd-1_v2c and the Col-0 gold standard sequence revealed a large inversion on chromosome 4 involving about 1 Mbp (Fig. 2). The left break point is at 1,637,889 bp and the right break point at 2,708,850 bp on chromosome 4 (NdChr4). The inverted sequence is 120 kbp shorter than the corresponding Col-0 sequence. PCR amplification of both inversion borders (AdditionalFile12) and Sanger sequencing of the generated amplicons was used to validate this rearrangement.

**Figure 2:**
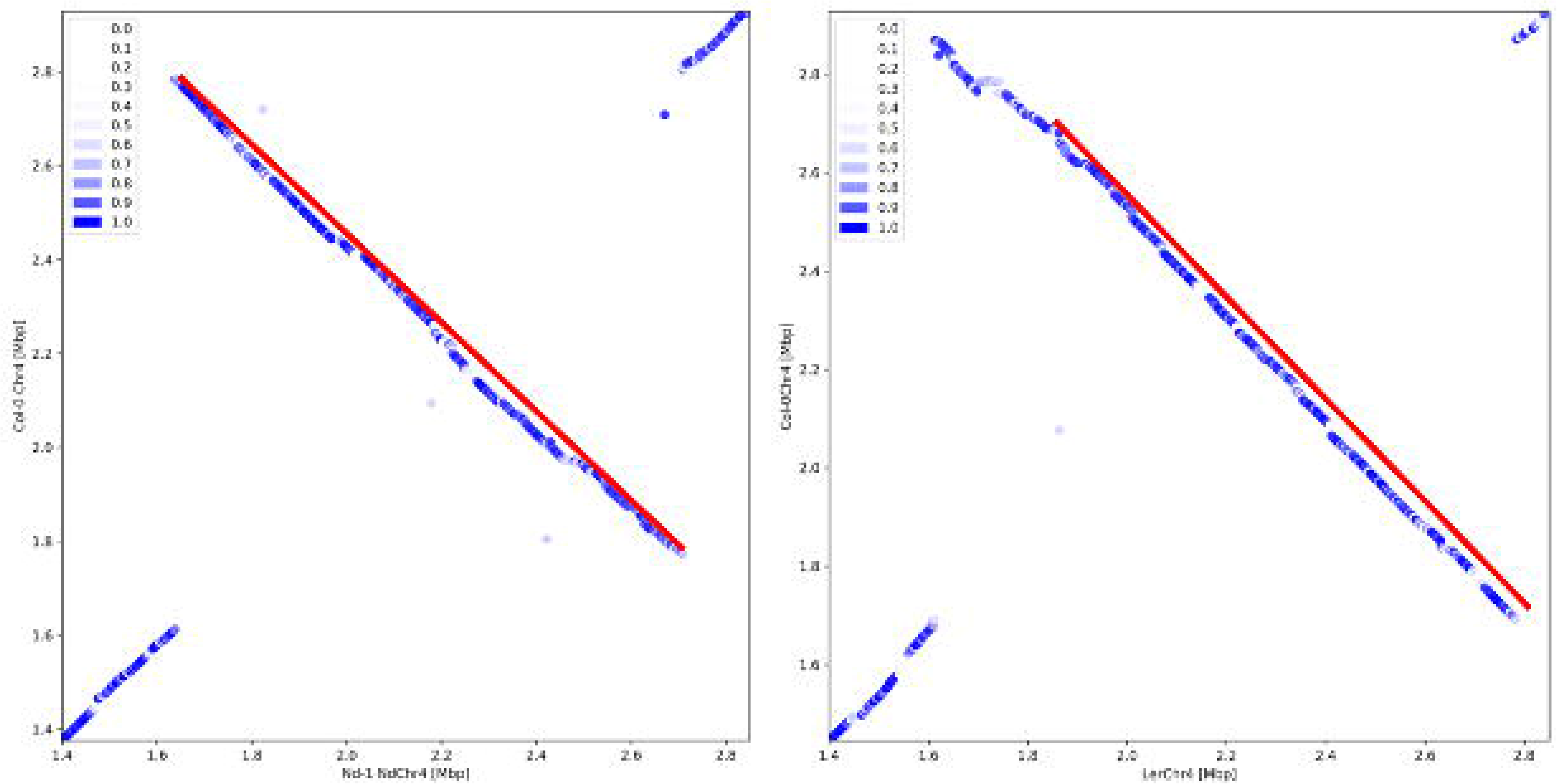
Inversion on chromosome 4. The dotplot heatmaps show the similarity between small fragments of two sequences. Each dot indicates a match of 1 kbp between both sequences, while the color is indicating the similarity of the matching sequences. A red line highlights an inversion between Nd-1 and Col-0 or L*er* and Col-0, respectively. A red arrow points at the position where the inversion alleles differ between Nd-1 and L*er*. (a) Comparison of the Nd-1 genome sequence against the Col-0 gold standard sequence reveals a 1 Mbp inversion. (b) The L*er* genome sequence displays another inversion allele [14].

The recombination frequency in this region was analyzed using the marker pair M84/M74. Only a single recombination was observed between these markers while investigating 60 plants. Moreover, only 8 recombination events in 108 plants were observed between another pair of markers, spanning a larger region across the inversion (AdditonalFile5). In contrast, the average recombination frequency per Mbp at the corresponding position on other chromosomes was between 12%, observed for M31/M32, and 18%, observed for M13/M14. Statistical analysis revealed a significant difference in the recombination frequencies between the corresponding positions on different chromosomes (p<0.001, prop.test() in R) supporting a reduced recombination rate across the inversion on chromosome 4.

Comparison of a region on Chr2, which is probably of mitochondrial origin (mtDNA), in the Col-0 gold standard sequence with the Nd-1 genome sequence revealed a 300 kbp highly divergent region (Fig. 3). Sequences between position 3.20 Mbp and 3.29 Mbp on NdChr2 of AthNd-1_v2c display low similarity to the Col-0 gold standard sequence, while there is almost no similarity between 3.29 Mbp and 3.48 Mbp. However, the length of both regions is roughly the same. Comparison against the L*er* genome sequence assembly revealed the absence of the entire region between 3.29 Mbp and 3.48 Mbp on chromosome 2. The Nd-1 sequence from this region lacks continuous similarity to any other region in the Col-0 or Nd-1 genome sequence. However, the 28 genes encoded in this region in Nd-1 show weak similarity to other Arabidopsis genes. Comparison of gene space sequences from this region against the entire Nd-1 assembly revealed some similarity on chromosome 3, 4, and 5 (AdditionalFile13).

**Figure 3:**
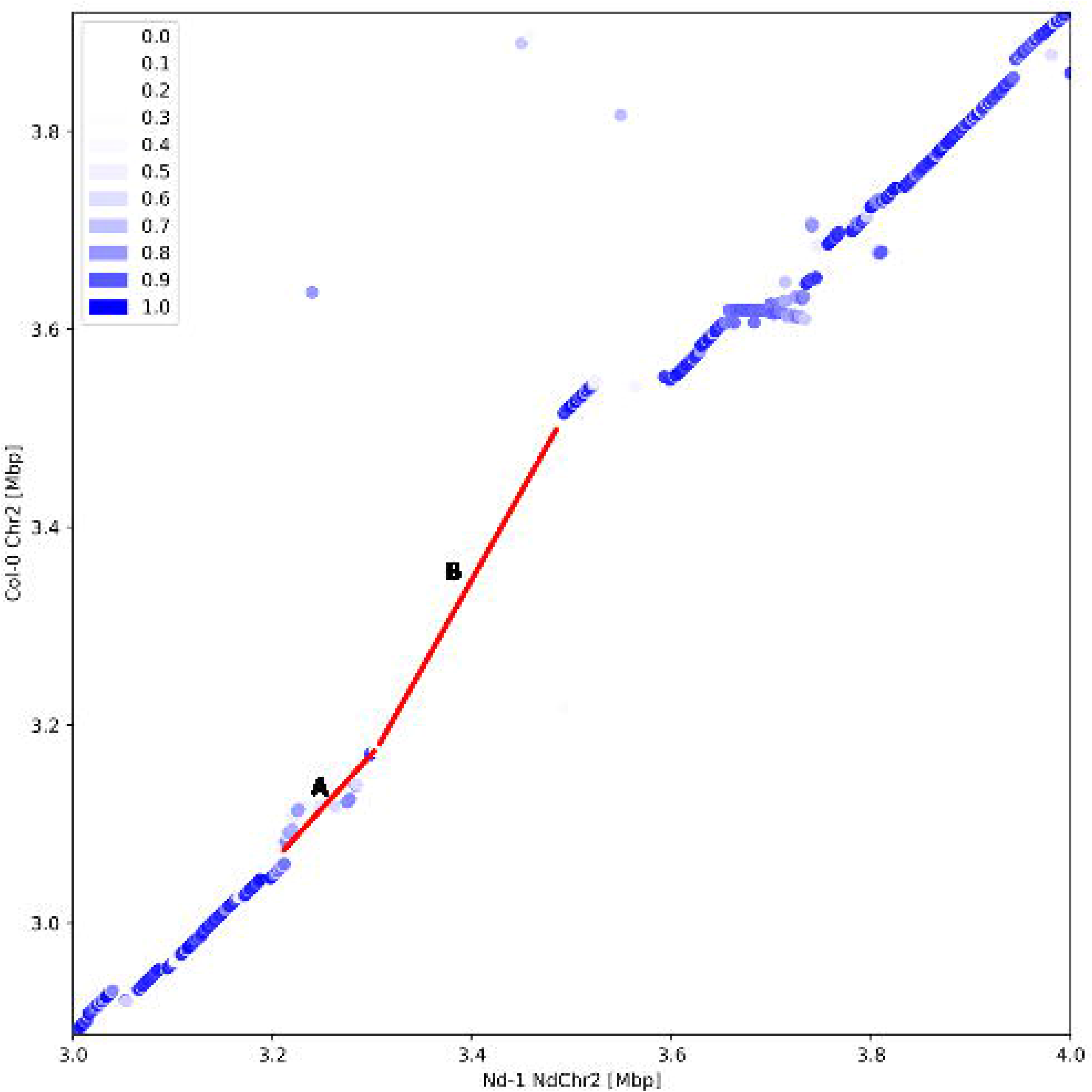
Highly divergent region on chromosome 2. There is a very low similarity region (light blue) between the sequences in region A and almost no similarity between the sequences in region B (white). The complete region between 3.29 Mbp and 3.48 Mbp on NdChr2 is missing in the L*er* genome.

An inversion on chromosome 3 of 170 kbp which was described between Col-0 and L*er* [14] is not present in Nd-1. The sequence similarity between Col-0 and Nd-1 is high in this region. In total, 2206 structural variants larger than 1 kbp were identified between Col-0 and Nd-1. The genome-wide distribution of these variants indicated a clustering around the centromeres (AdditionalFile14). A Spearman correlation coefficient of −0.79 (p=1.1*10^−27^) was calculated for the correlation of the number of SVs in a given interval and the distance of this interval to the centromere (AdditionalFile15). Therefore, these large structural variants are significantly more frequent in the centromeric and pericentromic regions. A total, of 148 new regions larger than 1 kbp were identified in Ath-Nd-1_v2c. These regions are also more frequent in proximity of the centromere (r=-0.43, p=3.7*10^−7^).

### Hint-based gene prediction

Hint-based gene prediction using AUGUSTUS with the *A. thaliana* species parameter set on the Nd-1 pseudochromosomes resulted in 30,126 nuclear protein coding genes (GeneSet_Nd-1_v2.0) with an average predicted transcript length of 1,798 bp, an average predicted CDS length of 1,391 bp and an average exon number per predicted transcript of 5.98. The number of predicted genes is reduced compared to the GeneSet_Nd-1_v1.1 [39] by 708 genes. In total, 28,042 (93%) representative peptide sequences were matched to Araport11 sequences and functionally annotated based on Araport11 information (AdditionalFile16). As controls we ran the gene prediction with same parameters on Col-0 and L*er* pseudochromosome sequences resulting in 30,352 genes and 29,302 genes, respectively. There were only minor differences concerning the average transcript and CDS length as well as the number of exons per gene.

Based on 35,636 TEs detected in Nd-1 (AdditionalFile8) 2,879 predicted Nd-1 genes were flagged as putative TE genes (AdditionalFile17, AdditionalFile18). This number matches well with the difference between the predicted genes in Nd-1 and the annotated protein coding genes in Araport11, which is supposed to be free of TE genes. The predicted mRNAs were supported by ESTs, which matched almost perfectly with an average similarity of 98.7% (AdditionalFile19). Additionally, the assembly was screened for TEs resulting in 613 consensus sequences of TE families (https://doi.org/10.5447/IPK/2019/4).

The comparative gene prediction with CAT resulted in 26,717 genes in Nd-1 and 26,681 genes in L*er*, respectively. Average CDS lengths were 1,292 bp and 1,291 bp, respectively. Since TE associated genes were not identified in this gene prediction process, the number of gene models predicted by CAT should be smaller than the number predicted by AUGUSTUS. The difference of 3,409 for Nd-1 is slightly exceeding the number of 2,879 TE associated genes that were predicted and flagged in the GeneSet_Nd-1_v2.0.

Besides protein encoding genes, 557 tRNA genes and 963 rRNA genes were predicted by INFERNAL. Comparison of these predicted tRNA genes to the result of tRNAscan-SE revealed an overlap of 552 (96.5%).

### Detection of gene space differences between Nd-1 and Col-0

A BLASTp-based comparison of all predicted Nd-1 peptide sequences and Col-0 Araport11 representative peptide sequences in both directions revealed 24,453 reciprocal best hits (RBHs) (AdditionalFile20). In total, 89.1% of all 27,445 nuclear Col-0 genes are represented in this RBH set. Analysis of the colinearity of the genomic location of all 24,453 RBHs (see AdditionalFile21 for a list) between Nd-1 and Col-0 showed overall synteny of both genomes as well as an inversion on chromosome 4 (AdditionalFile22). While most RBHs are properly flanked by their syntenic homologs and thus lead to a diagonal positioning of points in the scatter plot, there are 345 outliers (see AdditionalFile23 for a list). In general, outliers occur frequently in regions around the centromeres. Positional analysis revealed an involvement of many outliers in the large inversion on chromosome 4. An NGS read mapping at the positions of randomly selected outliers was manually inspected and indicated rearrangements between Nd-1 and Col-0. Structural variants, which affect at least three different genes in a row of RBH pairs, were identified from the RBH analysis. Examples beside the previously mentioned 1.2 Mbp inversion on chromosome 4 (*At4g03820-At4g05497*) are a translocation on chromosome 3 (*At3g60975-At3g61035*) as well as an inversion on chromosome 3 around *At3g30845*.

As a control we identified 25,556 (91.1%) RBHs between our gene prediction on Col-0 and the manually curated reference annotation Araport11. In addition, 24,329 (88.6 %) RBHs were identified between our gene prediction on the L*er* assembly and the Col-0 annotation Araport11.

In total, 947 protein encoding genes in Nd-1 (AdditionalFile24) were detected to be copies of only 421 genes annotated in Araport11. *SEC10* (*At5g12370*) [5] was previously described as an example for a tandem gene duplication collapsed in the Col-0 gold standard sequence. However, this region was already properly represented in AthNd-1_v1 [13]. Gene duplications of *At2g06555* (unknown protein), *At3g05530 (RPT5A*) and *At4g11510 (RALFL28*) in Nd-1 were confirmed by PCR amplification and Sanger sequencing of the sequences enclosed by both copies as well as through amplification of the entire event locus. On the other hand, there are 383 predicted genes in Nd-1 (AdditionalFile25) which appeared at least duplicated in Col-0.

### Pan-genomic analyses

Presence/absence variations of genes were inspected across a panel of 964 additional accessions (https://doi.org/10.5447/IPK/2019/4). In total, 25,809 genes were present in almost all accessions, 1,438 genes were considered dispensable, and the remaining 2,879 genes were flagged as TE or at least TE-associated genes (AdditionalFile26). Dispensable genes and TE genes are frequently located in proximity of the centromeres while core genes are less frequent in these regions (AdditionalFile27).

### Hidden locus in Col-0

*At4g22214* was identified as a gene duplicated in Nd-1 in our analysis. During experimental validation, we did not detect the expected difference between Col-0 and Nd-1 DNA concerning the locus around *At4g22214*. However, the PCR results from Col-0 matched the expectation based on the Nd-1 genome sequence thus suggesting a collapsed gene sequence in the Col-0 gold standard sequence. This hypothesis was supported by PCR results with outwards facing primers (Fig. 4). Cloning of the At4g22214 region of Col-0 in five overlapping fragments was done to enable Sanger sequencing. The combination of Sanger and paired-end Illumina sequencing reads revealed a tandem duplication with modification of the original gene (Fig. 4). The copies were designated At4g22214a and At4g22214b based on their position in the genome (GenBank: MG720229). While At4g22214b almost perfectly matches the Araport11 annotation of At4g22214, a significant part of the CDS of At4g22214a is missing. Therefore, the gene product of this copy is probably functionless. At4g22214 is annotated as defensin-like family protein [55]. Since the family of defensin proteins contains about 300 members in *A. thaliana* [55], the functional implications of this duplication are probably low.

**Figure 4:**
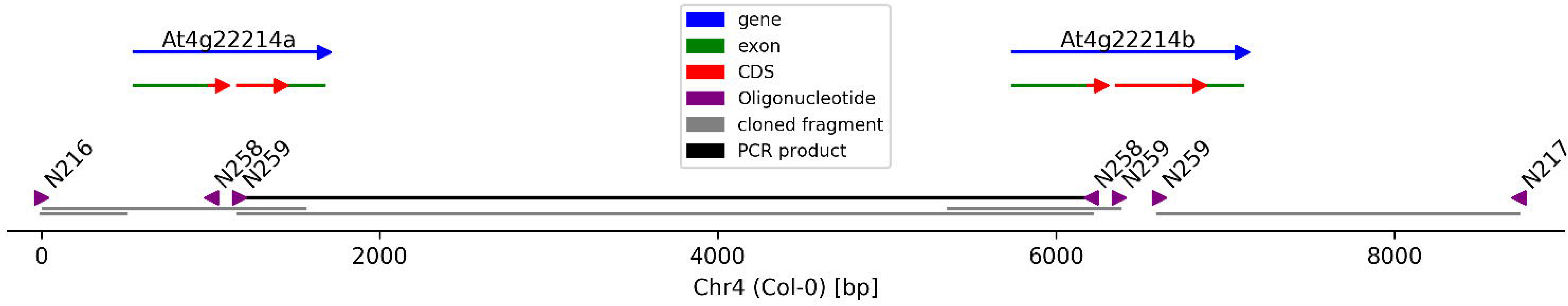
Hidden locus in the Col-0 reference sequence. Differences between the Nd-1 and Col-0 genome sequences lead to the discovery of a collapsed region in the Col-0 gold standard sequence. There are two copies of At4g22214 (blue) present in the Col-0 genome, while only one copy is represented in the Col-0 gold standard sequence. This gene duplication was initially validated through PCR with outwards facing oligonucleotides N258 and N259 (purple) which lead to the formation of the expected PCR product (black). Parts of this region were cloned into plasmids (grey) for sequencing. Sanger and paired-end Illumina sequencing reads revealed one complete gene (At4g22214b) and a degenerated copy (At4g22214a). Moreover, the region downstream of the complete gene copy in Nd-1 indicates the presence of at least one additional degenerated copy.

### Gaps in the Col-0 reference sequence

Despite its very high quality, the Col-0 gold standard sequence contains 92 N stretches of various sizes representing regions of unknown sequence like the NOR clusters or centromeres. Ath-Nd1_v2c enabled the investigation of some of these sequences based on homology assumptions. A total of 13 Col-0 gold standard sequence gaps were spanned with high confidence by Ath-Nd1_v2c and therefore selected for homopolymer frequency analysis. The corresponding regions in Nd-1 are significantly enriched with homopolymers in comparison to randomly picked control sequences (p=0.000048, Mann-Whitney U test) (AdditionalFile28).

## Discussion

### Genome structure of the *A. thaliana* accession Nd-1

In order to further investigate large variations in the range of several kbp up to several Mbp between *A. thaliana* accessions, we performed a *de novo* genome assembly for the Nd-1 accession using long sequencing reads and cutting-edge assembly software. Based on SMRT sequencing reads the assembly contiguity was improved by over 75 fold considering the number of contigs in the previously released NGS-based assembly [13]. Assembly statistics were comparable to other projects using similar data [11, 14, 15, 56, 57]. Despite the very high contiguity, regions like NORs still pose a major challenge. These regions are not just randomly clustered repeats, but highly controlled and ordered repetitions of sequences [58]. Therefore, the identification of accession-specific sequence differences could explain phenotypic differences. NOR repeat unit sequences in Ath-Nd-1_v2c are located at on the distal short arm of NdChr2 and NdChr4. This NOR position matches the situation in Col-0 where the NOR2 is located distal on the short arm of Chr2 [59] and NOR4 on Chr4, respectively. In addition to NORs, the assembly of chromosome ends remains still challenging, since the absence of some telomeric sequences in a high quality assembly was observed before [14]. Despite the absence of challenging repeats, regions close to the telomeres including the genes At3g01060 (BUSCO ID: EOG09360D4T) and At5g01010 (BUSCO ID: EOG09360DFK) were not assembled by FALCON (Ath-Nd1_v2f) although sequence reads covering these regions were present in the input data. Also, this region is represented in the Canu assembly (Ath-Nd1_v2c). In our hands, Canu performed best for the assembly of the Nd-1 genome sequence based on SMRT sequencing reads.

### Nuclear genome sequence differences

The increased contiguity of this long read assembly was necessary to discover an approximately 1 Mbp inversion through sequence comparison as well as RBH analysis. An earlier Illumina short read based assembly [13] lacked sufficient contiguity in the region of interest to reveal both breakpoints of this variant between Col-0 and Nd-1. The large inversion at the north of NdChr4 relative to the Col-0 gold standard sequence turned out as a modification of the allele originally detected in L*er* [14, 60]. However, the Nd-1 allele is different from the L*er* allele. This could explain previous observations in several hundred *A. thaliana* accessions, which share the left inversion border with L*er*, but show a different right inversion border [14].

Despite the very much improved contiguity and selection of Canu as the currently best assembler for our dataset, there are only very small parts of pericentromeric sequences represented in the Ath-Nd1_v2c assembly. Centromeric regions with an estimated size of 5 Mbp each [13] were not assembled. The absence of these highly repetitive regions is in agreement with previous findings that the error rate of long reads is still too high to resolve NORs and centromeres [15]. The pericentromeric and to a very limited extent also centromeric regions are represented in the assemblies by small contigs of 50-100 kbp. However, absence of telomeric sequences from some pseudochromsome ends was observed before even for a very high quality assembly [14, 15]. The detected telomeric repeats at the centromere positions support previously reported hypothesis about the evolution of centromers out of telomere sequences [61], and telomeric or centromeric sequences, respectively, at the end of at least some pseudochromosomes indicated the completeness of the Ath-Nd1_v2c assembly at these points. However, almost 20 years after the release of the first chromosome sequences of *A. thaliana*, we are still not able to assemble complete centromere sequences continuously.

Sequence differences observed on the short arm of chromosome 2 between Col-0 and Nd-1 could be due to the integration of mtDNA into the Chr2 of Col-0 [2]. This region was reported to be collapsed in the Col-0 reference genome sequence, harboring in real Col-0 DNA about 600 kbp of mtDNA instead of the 270 kbp represented in the reference genome sequence [62]. However, Ath-Nd1_v2c did not reveal such a duplication of the mtDNA located in NdChr2. Filtering of plastome and chondrome sequences during the polishing process could be responsible for this if copies would have been fragmented into multiple contigs. In this region, we detected in the Nd-1 assembly sequences unrelated to the corresponding Col-0 reference sequence. Since Nd-1 genes of this region show similarity to gene clusters on other chromosomes, they could be relics of a whole genome duplication as reported before for several regions of the Col-0 gold standard sequence [63]. This difference on Chr2 at about 3.2 to 3.5 Mbp is only one example for a large variant region between Col-0 and Nd-1. Similar though shorter such differences detected around centromeres could be explained by TEs and pseudogenes which were previously reported as causes for intra-species variants in these regions [62, 64].

### Plastome and chondrome

Size and structure of the Nd-1 plastome is very similar to Col-0 [2] or L*er* [27]. In accordance with the overall genome similarities, the observed number of small differences between the plastome sequences of Col-0 and Nd-1 is slightly higher than the value reported before for the Col-0 comparison to L*er* [27].

The size of the Nd-1 chondrome matches previously reported values for the large chondrome configuration of other *A. thaliana* accessions [65]. Large structural differences between the Col-0 chondrome [65] and the Nd-1 chondrome could be due to the previously described high diversity of this subgenome including the generation of substoichiometric DNA molecules [66, 67]. In addition, the mtDNA level was reported to differ between cell types or cells of different ages within the same plant [68, 69]. The almost equal read coverage of the assembled Nd-1 chondrome could be explained by the young age of the plants at the point of DNA isolation, as the amount of all chondrome parts should be the same in young leafs [69].

### Nd-1 gene space

Many diploid plant genomes contain on average around or even slightly below 30,000 protein encoding genes [35, 70] with the *A. thaliana* genome harboring 27,445 nuclear protein-coding genes according to the most recent Araport11 annotation [40]. Since there are only few other chromosome-level assembly sequences of *A. thaliana* available at the moment, we do not know the precise variation range of gene numbers between different accessions. The number of 27,247 predicted non-TE genes in Nd-1 is further supported by the identification of 24,453 RBHs with the Araport11 [40] annotation of the Col-0 gold standard sequence. This number exceeds the matches between Col-0 and Ler-0 [14]. Our chromosome-level assembly further enhances the gene prediction quality since at least 89.1% of all Col-0 genes were recovered. This reinforces previous studies that also reported annotation improvements through improved assembly quality [71]. The number of 35,636 TEs annotated for Ath-Nd-1_v2c exceeds the number of 33,892 such elements identified in the previous assembly version Ath-Nd-1_v1 [13] by 1,744. Such an increase in the number of resolved TEs as well as an improved assembly of TE-rich regions in assemblies based on long reads was reported before [72].

Due to the high proportion of genes within the *A. thaliana* genome assigned to paralogous groups with high sequence similarity [73, 74], we speculated that the identification of orthologous pairs via RBH analysis might be almost saturated. Gene prediction with the same parameters on the Col-0 gold standard genome sequence prior to RBH analysis supported this hypothesis. However, the incorporation of accession-specific RNA-Seq derived hints could further increase the accuracy of the Nd-1 gene prediction. Since there are even some RBHs at non-syntenic positions between the control Col-0 annotation and the Araport11 annotation, our Nd-1 annotation is already of very high accuracy. The precise annotation of non-canonical splice sites via hints as described before [39] contributed to the new GeneSet_Nd-1_v2.0. Gene duplication and deletion numbers, or accession specific presence/absence variations of genes, in Nd-1 and Col-0 are in the same range as previously reported values of up to a few hundred [14, 75]. Since we were searching genome-wide for copies of genes without requiring an annotated feature in each genome sequence, both numbers might include some pseudogenes due to the frequent occurrence of these elements within plant genomes [76, 77]. Since all comparisons rely on the constructed sequences we cannot absolutely exclude that a small number of other genes were detected as amplified due to collapsed sequences similar to SEC10 (At5g12370) [5]. Removing TE genes based on sequence similarity to annotated features reduced the proportion of putative pseudogenes. However, it is impossible to unequivocally distinguish between real genes and pseudogenes in all cases, because even genes with a premature stop codon or a frameshift mutation could function as a truncated version or give rise to regulatory RNAs [74, 77-79]. In addition, the impact of copy number variations involving protein encoding genes in *A. thaliana* might be higher than previously assumed thus supporting the existence of multiple gene copies [80]. Gene expression analysis could support the discrimination of pseudogenes, because low gene expression in *A. thaliana* was reported to be associated with pseudogenization [81]. Despite the unclear status of the gene product, the mere presence of these sequences revealed fascinating insights into genome evolution and contributed to the pan-genome [82, 83].

## Conclusions

We report a high quality long read *de novo* assembly (AthNd-1_v2c) of the *A. thaliana* accession Nd-1, which improved significantly on the previously released NGS assembly sequence AthNd-1_v1.0 [13]. Comparison of the GeneSet_Nd-1_v2.0 with the Araport11 nuclear protein coding genes revealed 24,453 RBHs supporting an overall synteny between both *A. thaliana* accessions except for an approximately 1 Mbp inversion at the north of chromosome 4. Moreover, large structural variants were identified in the pericentromeric regions. Comparisons with the Col-0 gold standard sequence also revealed a collapsed locus around At4g22214 in Col-0. Therefore, this work contributes to the increasing *A. thaliana* pan-genome with significantly extended details about genomic rearrangements.

## Supporting information

AdditionalFile1

AdditionalFile2

AdditionalFile3

AdditionalFile4

AdditionalFile5

AdditionalFile6

AdditionalFile7

AdditionalFile8

AdditionalFile9

AdditionalFile10

AdditionalFile11

AdditionalFile12

AdditionalFile13

AdditionalFile14

AdditionalFile15

AdditionalFile16

AdditionalFile17

AdditionalFile18

AdditionalFile19

AdditionalFile20

AdditionalFile21

AdditionalFile22

AdditionalFile23

AdditionalFile24

AdditionalFile25

AdditionalFile26

AdditionalFile27

AdditionalFile28

## List of abbreviations

NGS: next generation sequencing
NOR: nucleolus organizing region
RBH: reciprocal best hit
SMRT: single molecule real time

## Declarations

### Ethics approval and consent to participate

Not applicable

### Consent for publication

Not applicable

### Availability of data and materials

The data sets supporting the results of this article are included within the article and its additional files. The Ath-Nd-1_v2 assemblies (Table 1) and derived files are available as external downloads from http://doi.org/10.5447/IPK/2019/4. Sequencing reads were submitted to the SRA (SRP066294). Python scripts developed and applied for this study are available on github: https://github.com/bpucker/Nd1_PacBio (http://doi.org/10.5281/zenodo.2590750).

### Author’s contributions

BP, DH and BW conceived and designed research. BP, KS, KF, BH and RR conducted experiments. BP, DH and BW interpreted the data. BP and BW wrote the manuscript. All authors read and approved the final manuscript.

## Acknowledgements

We thank Willy Keller for isolating high molecular DNA, Katharina Kemmet for extensive genotyping of plants for the genetic map, Helene Schellenberg, Ann-Christin Polikeit and Prisca Viehöver for Sanger sequencing, and Melanie Kuhlmann as well as Andrea Voigt for taking excellent care of the plants. We are very grateful for support from the Bioinformatics Resource Facility and Stefan Albaum.

## Additional Files

**AdditionalFile1. Sequencing Statistics**.

Statistical information about the generated SMRT sequencing data for the *A. thaliana* Nd-1 genome assembly are listed in this table. The expected genome size is based on several analyses reporting values around 150 Mbp [84, 85].

**AdditionalFile2. Canu assembly parameters**.

Listing of all parameters that were adjusted for the Canu assembly of the Nd-1 nucleome. While most default parameters were kept, some were specifically adjusted for this plant genome assembly.

**AdditionalFile3. FALCON assembly parameters**.

All parameters that were adjusted for the FALCON assembly of the Nd-1 nucleome are listed in this table. While most default parameters were kept, some were specifically adjusted for this plant genome assembly.

**AdditionalFile4. Molecular markers for genetic linkage analysis**.

All markers require the amplification of a genomic region using the listed oligonucleotides under the specified conditions (annealing temperature, elongation time). Depending on the fragment size differences, the resulting PCR products can allow the separation of both alleles by agarose gel electrophoresis (length polymorphism) or might require Sanger sequencing to investigate single SNPs.

**AdditionalFile5. Distribution of genetic markers over physical map**.

The positions of all genetic markers on the pseudochromosome sequences are illustrated. Assembled sequences were positioned based on the genetic linkage information. Some genetic marker combinations allowed the investigation of recombination frequencies within continuous sequences.

**AdditionaiFile6. Oligonucleotide sequences for genetic linkage analysis**.

Sequences, names and recommended annealing temperatures of all oligonucleotides used in this work are listed in this table. Usage remarks for the oligonucleotides are provided as well.

**AdditionalFile7. Alignment of Ath-Nd1_v2c and Ath-Nd1_v2f**.

Assemblies generated by Canu and FALCON, respectively, were compared via BLASTn search of 10 kb sequence chunks. Color and position of dots in the figure indicate the position of the best hit on the respective sequence.

**AdditionalFile8. TE positions in the Nd-1 genome sequence**.

TE genes, TEs, and TE fragments in the Nd-1 genome sequence were identified based on sequence similarity to annotated TEs from the Col-0 gold standard sequence (Araport11) [40].

**AdditionalFile9. BUSCO analysis of the Col-0 and Nd-1 genome sequences**.

BUSCO v2.0 was run on the genomic sequences of Col-0 and Nd-1 using AUGUSTUS 3.3 with default parameters for the gene prediction process.

**AdditionalFile10. Nd-1 plastome map**.

The GC content (black) and GC skew (green for positive GC skew, purple for negative GC skew) of the plastome sequence were analyzed by CGView [31]. The sequence and its properties are very similar to the Col-0 plastome sequence.

**AdditionalFile11. Nd-1 chondrome map**.

The GC content (black) and GC skew (green for positive GC skew, purple for negative GC skew) of the chondrome sequence were analyzed by CGView [31]. The sequence and its properties are very similar to the Col-0 chondrome sequence.

**AdditionalFile12. Experimental validation of 1 Mbp inversion on chromosome 4**.

The identified inversion between Nd-1 and Col-0 on chromosome 4 is different from the inversion described before between Col-0 and L*er* [14]. However, the left breakpoint is the same for both alleles enabling the use of previously published oligonucleotide sequences [14]. The right breakpoint was identified by manual investigation of sequence alignments. Both breakpoints were validated via PCR using the oligonucleotides (for sequences see AdditionalFile6) as illustrated in (a). The results support the expected inversion borders (b).

**AdditionalFile13. Genome-wide distribution of genes inserted on chromosome 2 in Nd-1**.

AthNd-1_v2c and the Col-0 gold standard sequence display a highly diverged region at the north of chromosome 2, which is about 300 kbp long. BLASTn of the complete Nd-1 gene sequences from this region revealed several regions on other Nd-1 chromosomes with copies of these genes.

**AdditionalFile14. Genome-wide distribution of large structural variants**.

The distribution of structural variants (SVs) >10 kbp (red dots) between Col-0 and Nd-1 over all five pseudochromosome sequences (black lines) is illustrated. Additionally, the assumed centromere (CEN) positions are indicated (blue dots). Most SVs are clustered in the (peri-)centromeric region.

**AdditionalFile15. Clustering of SVs around centromeres**.

The correlation between the number of SVs in a given part of the genome sequence (1 Mbp) and the distance of this region to the centromere position is illustrated. SVs are clustered around the centromeres (Spearman correlation coefficient = −0.66, p-value = 1.7*10^−16^).

**AdditionalFile16. Functional annotation of GeneSet_Nd-1_v2.0**.

Functional annotations were transferred from Araport11 to corresponding RBHs in GeneSet_Nd-1_v2.0. In addition, genes were annotated based on the best BLAST hit if annotation via RBH was not possible.

**AdditionalFile17. TE overlap with GeneSet_Nd-1_v2.0**.

The overlap between annotated TEs (AdditionalFile8) and predicted protein coding genes was analyzed to identify TE genes. This figure illustrates the fraction of a gene that is covered by a TE. Since TEs might occur within the intron of a gene, only genes with at least 80% TE coverage were flagged as TE genes (AdditionalFile18).

**AdditionalFile18. TE genes in GeneSet_Nd-1_v2.0**.

These genes were predicted by AUGUSTUS as protein coding genes. Due to their positional overlap with TEs (AdditionalFile8), they were flagged as TE genes and excluded from further gene set analysis.

**AdditionalFile19. EST mapping**.

Percentage of nucleotides in ESTs matching predicted transcripts are displayed.

**AdditionalFile20. Gene set overlap between Araport11, GeneSet_Nd-1_v1.1, and GeneSet_Nd-1_v2.0**.

RBHs were identified pairwise between gene sets. The overlap was identified by mapping all genes onto Araport11 identifiers. Venn diagram construction was performed at http://bioinformatics.psb.ugent.be/webtools/Venn/.

**AdditionalFile21. Reciprocal best hits (RBH) pairs between Col-0 and Nd-1**.

Reciprocal best hits between predicted peptide sequences of Nd-1 and the representative peptide sequences of Col-0 (Araport11).

**AdditionalFile22. Reciprocal best hits (RHB) indicates inversion between Nd-1 and Col-0**.

Genes in RBH pairs were sorted based on their position on the five pseudochromosomes of the two genome sequences to form the x (Col-0) and y (Nd-1) axes of this diagram. Plotting the positions of each RBH pair leads to a bisecting line of black dots representing genes at perfectly syntenic positions. Red and green dots indicate RBH gene pair positions deviating from the syntenic position. Red dots symbolize a unique match to another gene, while green dots indicate multiple very similar matches. Positions of the centromere (CEN4) on the chromosomes of both accessions are indicated by purple lines. An inversion involving 131 genes in RBH pairs just north of CEN4 distinguishes Nd-1 and Col-0.

**AdditionalFile23. RBH outliers in GeneSet_Nd-1_v2.0**.

RBHs between Araport11 and GeneSet_Nd-1_v2.0 were identified based on encoded representative peptide sequences. All 242 RBHs at positions deviating from the syntenic diagonal line were collected. The functional annotation of these genes was derived from Araport11.

**AdditionalFile24. Duplicated genes in Nd-1**.

The listed 385 Col-0 genes (Araport11 [40]) have at least two copies in Nd-1. Exons of these genes showed an increased copy number in Ath-Nd-1_v2c compared to the Col-0 gold standard sequence. The annotation was derived from Araport11.

**AdditionalFile25. Duplicated genes in Col-0**.

The listed 394 Nd-1 genes have at least two copies in Col-0. Exons of these genes showed an increased copy number in the Col-0 gold standard sequence compared to Ath-Nd-1_v2c.

**AdditionalFile26. Classification of genes as core, dispensable, or TE**.

GeneSet_Nd-1_v2.0 genes were classified as core, dispensable, or TE genes based on coverage in a read mapping involving 1,137 Illumina read data sets.

**AdditionalFile27. Genome-wide distribution of core, dispensable, and TE genes**.

Visualisation of the position of core genes, dispensable genes and TEs along the chromosomes.

**AdditionalFile28. Critical regions in the Col-0 gold standard sequence**.

The high contiguity of the Ath-Nd-1_v2c assembly enabled the investigation of 13 sequences corresponding to gaps in the Col-0 gold standard sequence. This figure illustrates the homotetranucleotide occurrence in these sequences (red dots) in comparison to some randomly selected reference sequences (green dots). While there is a clear enrichment of homotetranucleotides in the gap-homolog sequences, there was no clear correlation between the length of a gap and the composition of the corresponding sequence observed.

