## Supplementary figures and images for "A Chromosome-level Sequence Assembly Reveals the Structure of the *Arabidopsis thaliana* Nd-1 Genome and its Gene Set"

### AdditionalFile5

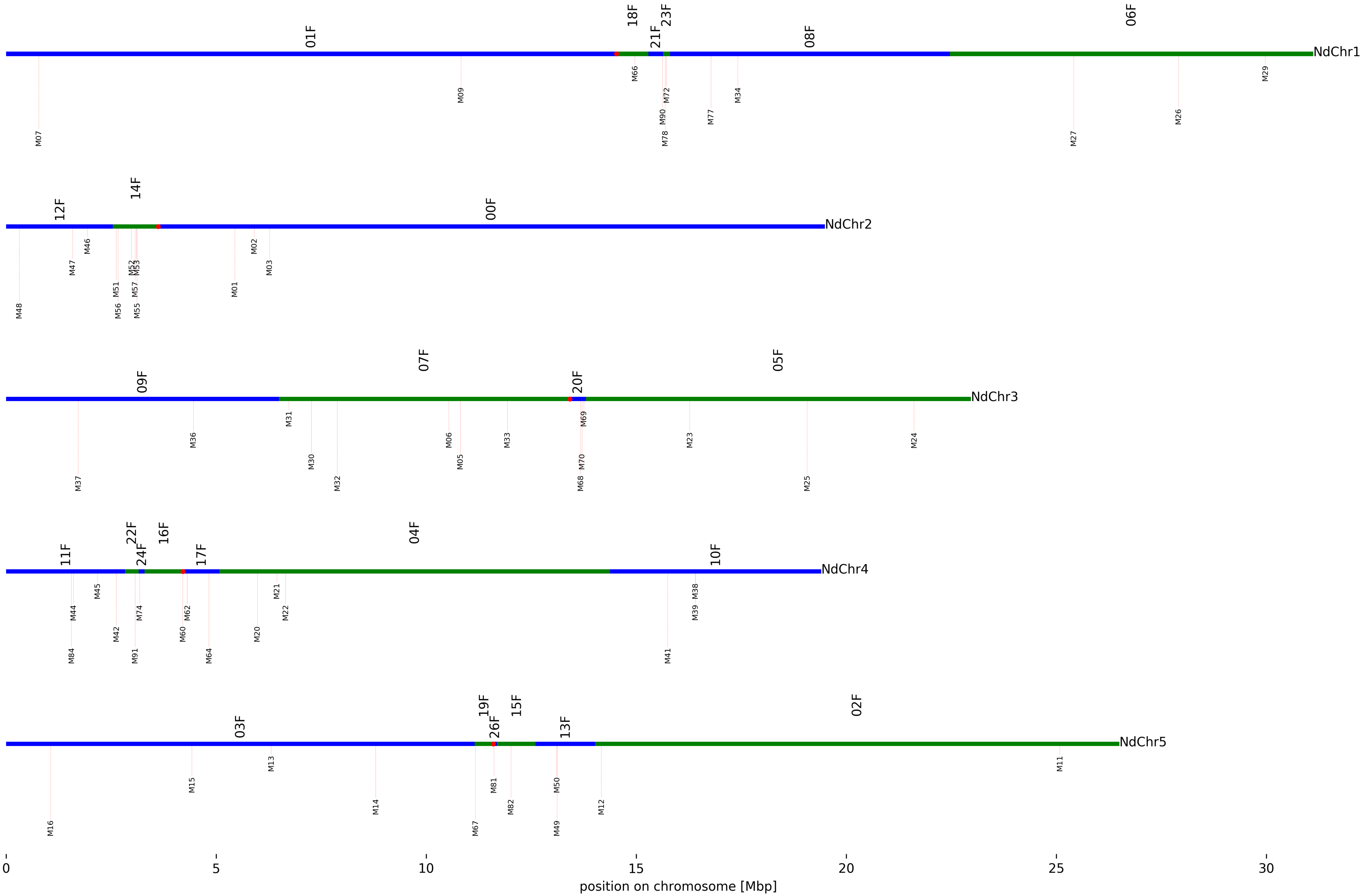

### AdditionalFile7

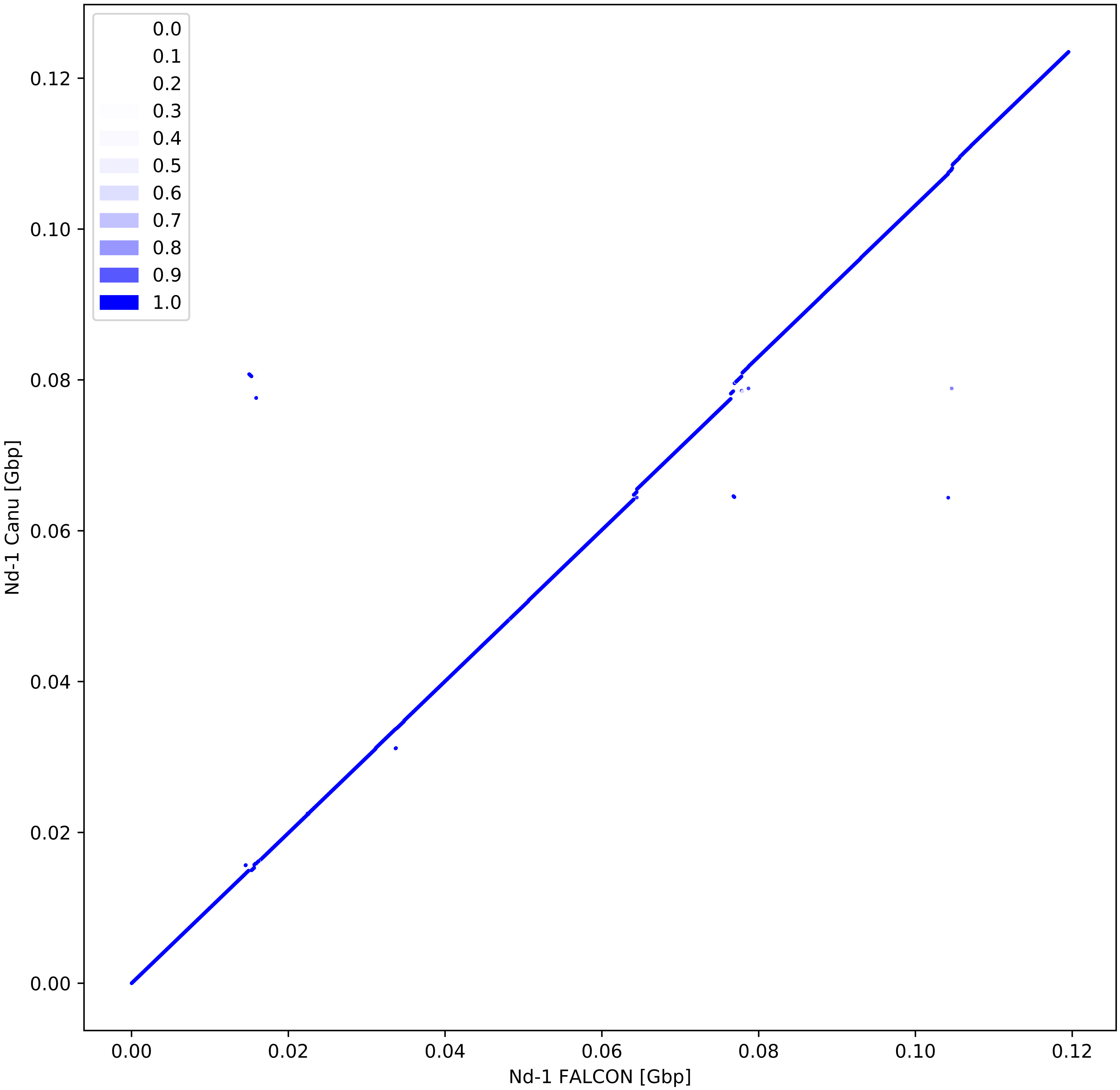

### AdditionalFile10

Length: 154,443 bp

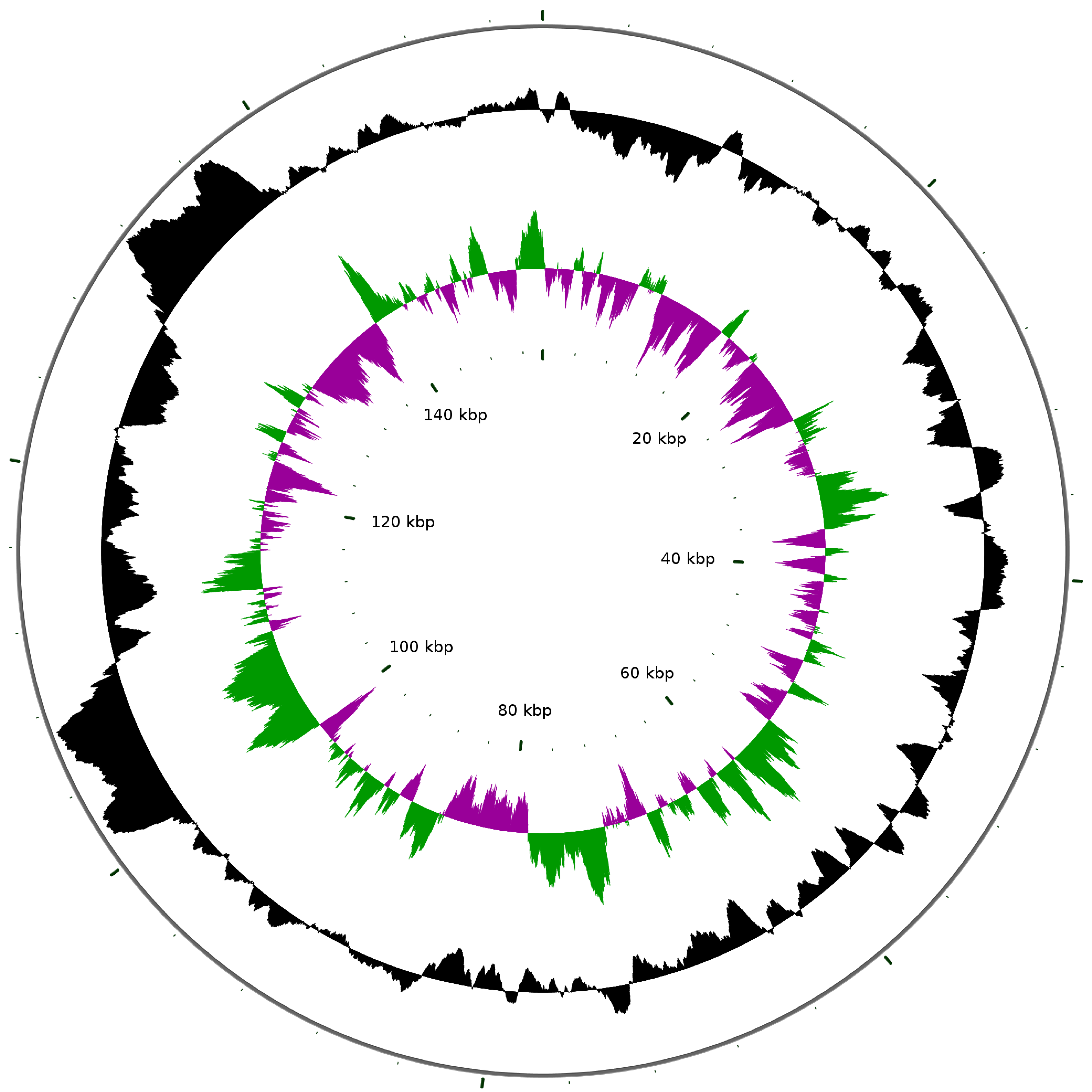

### AdditionalFile11

Length: 368,216 bp

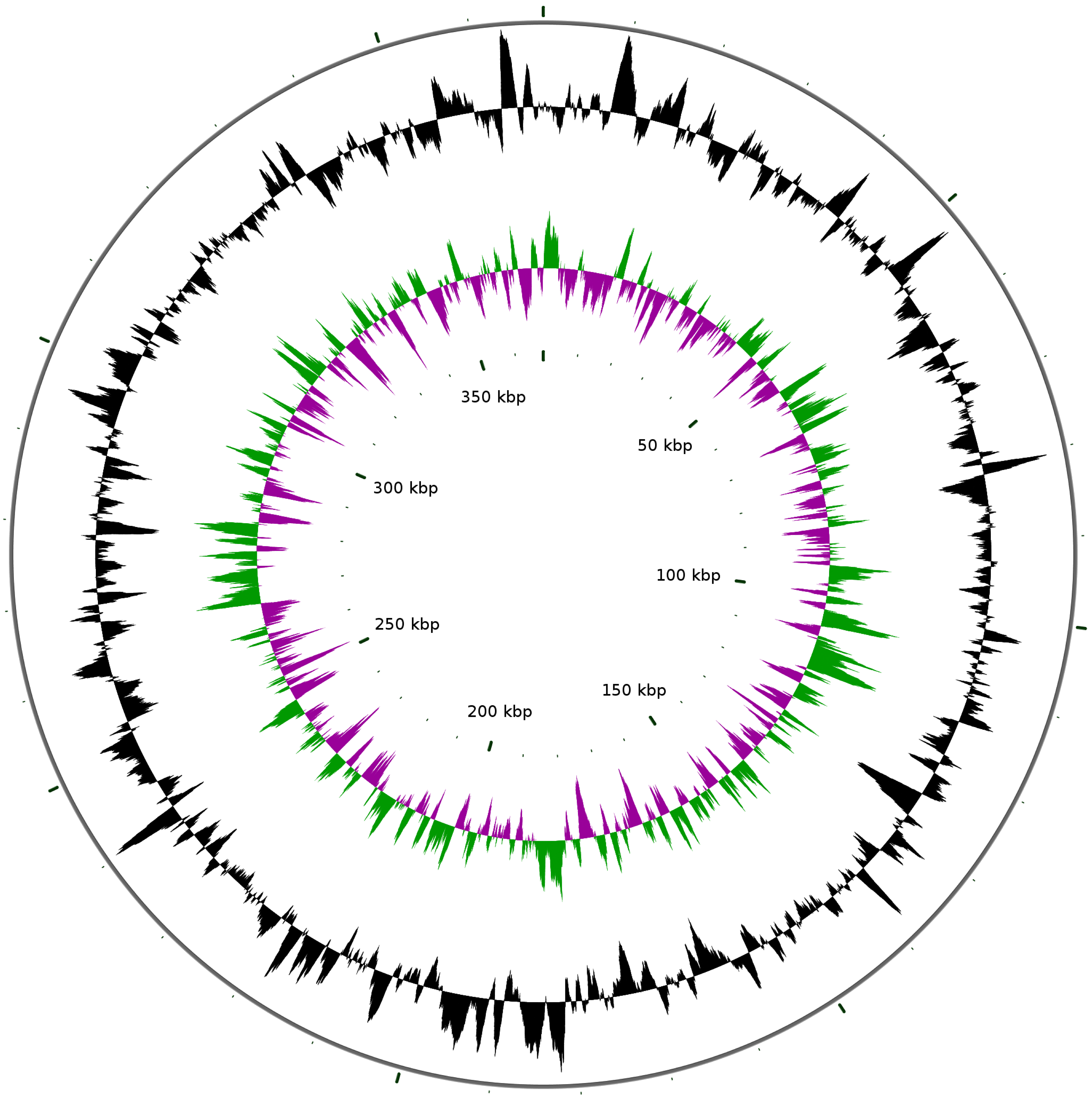

### AdditionalFile12

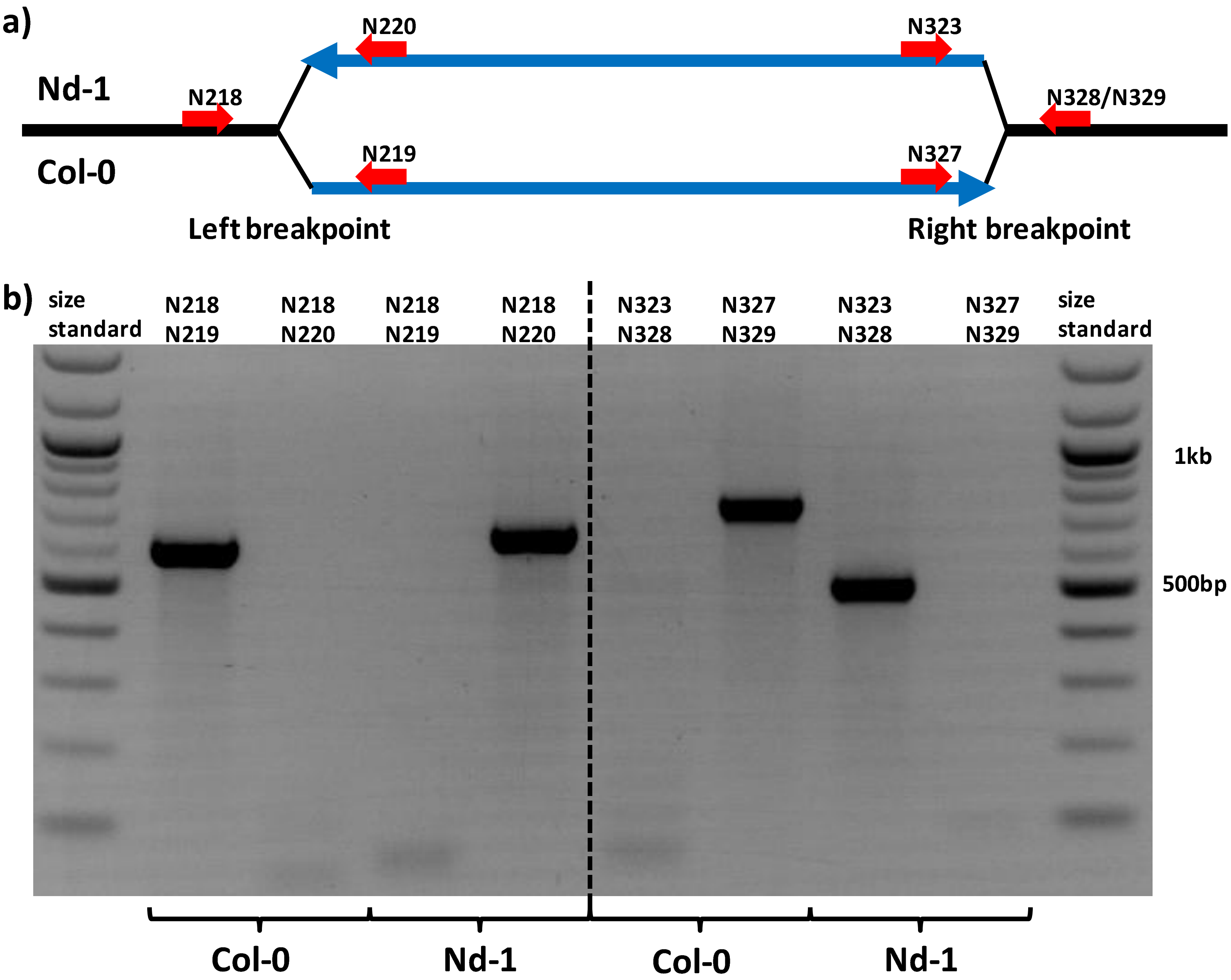

### AdditionalFile13

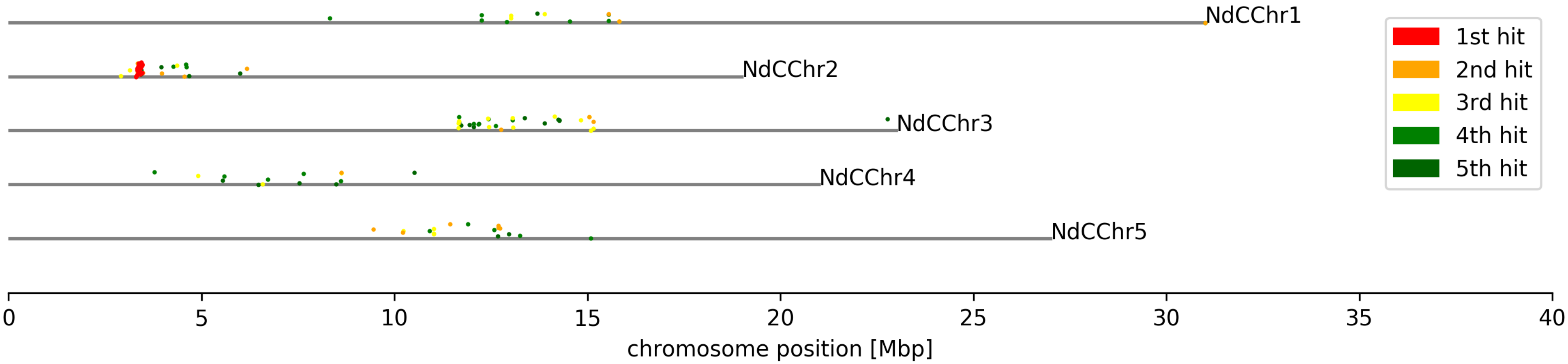

### AdditionalFile14

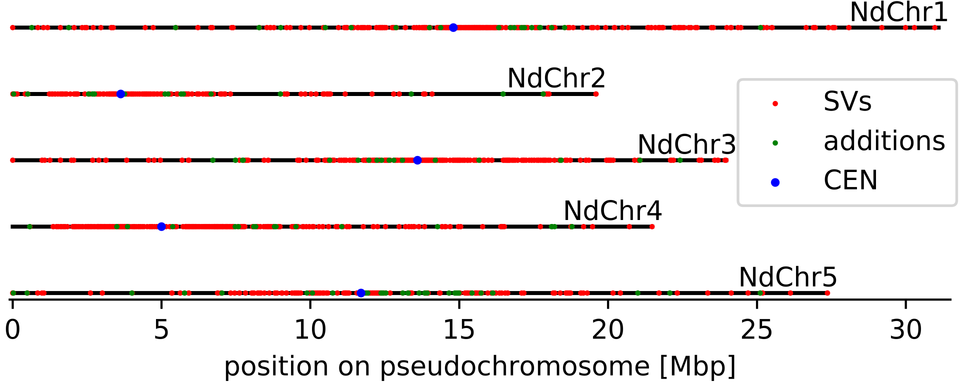

### AdditionalFile15

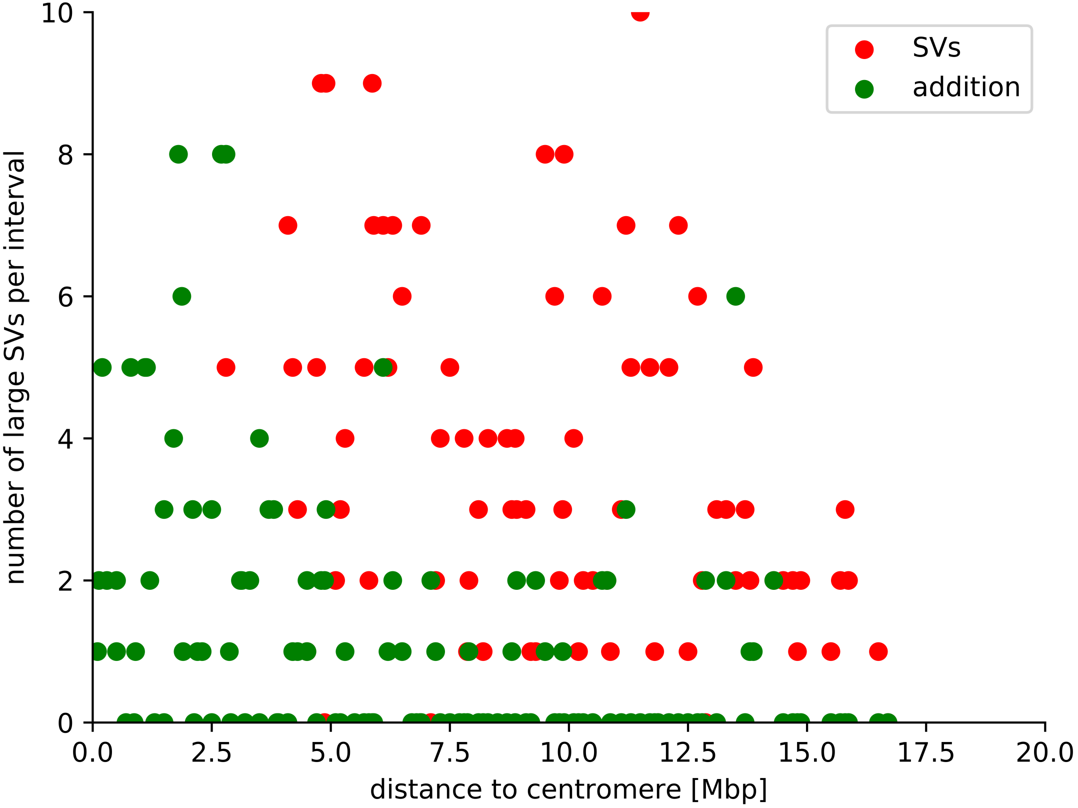

### AdditionalFile17

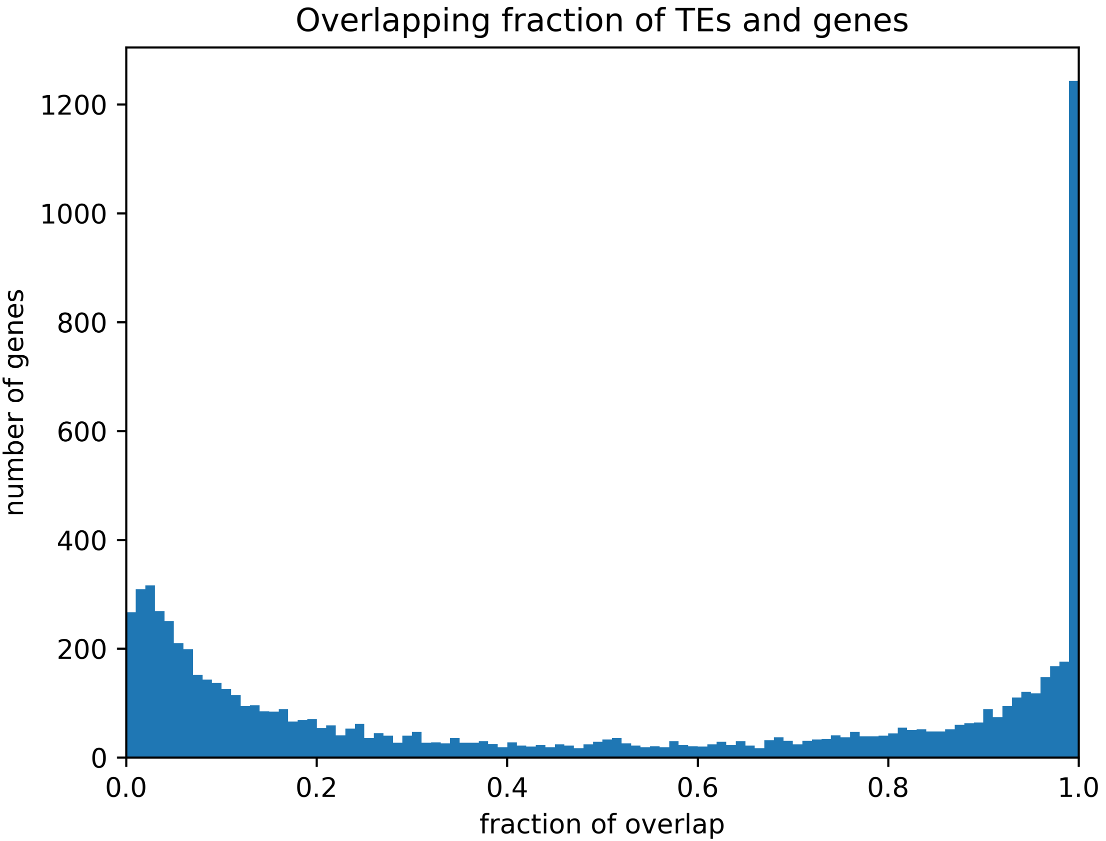

### AdditionalFile19

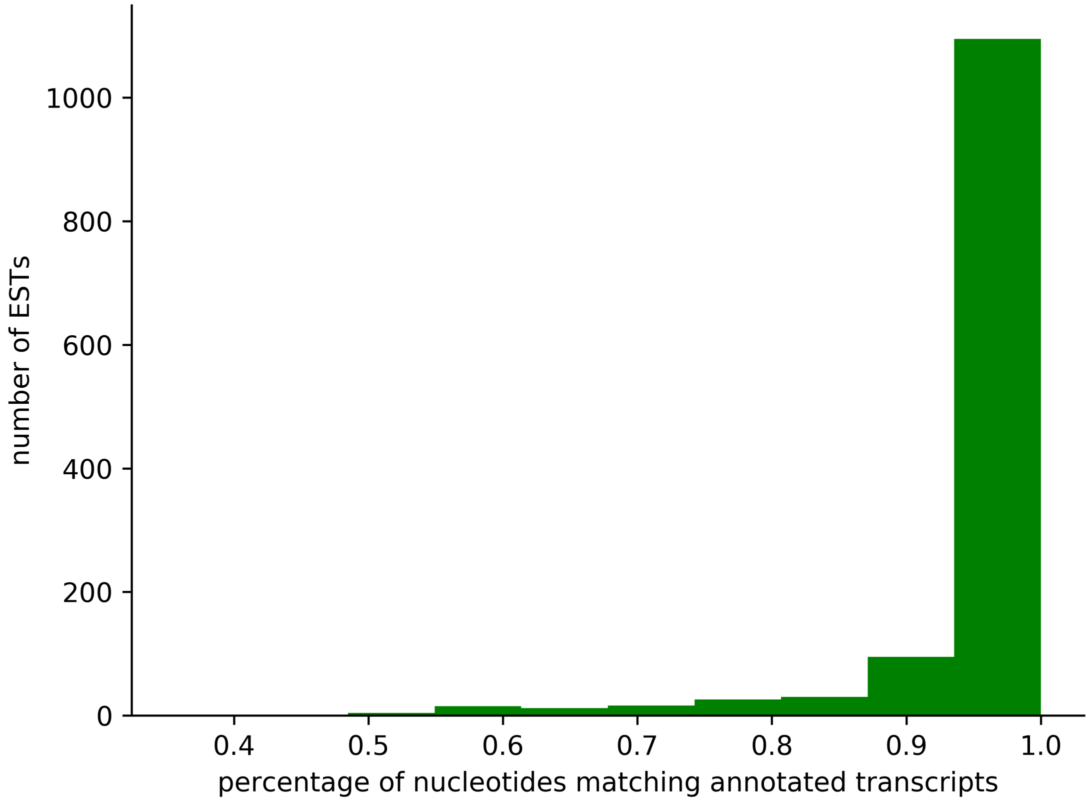

### AdditionalFile20

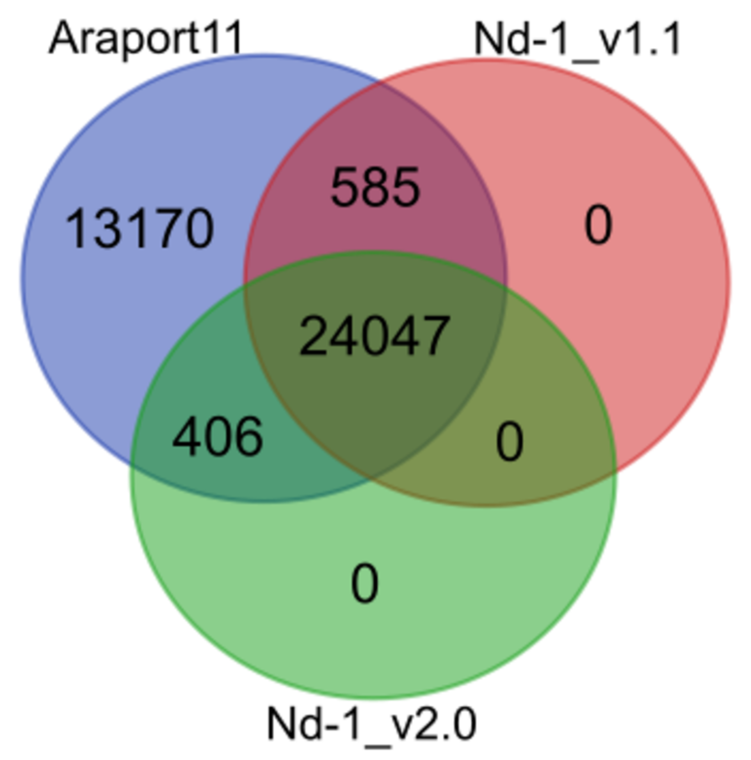

### AdditionalFile22

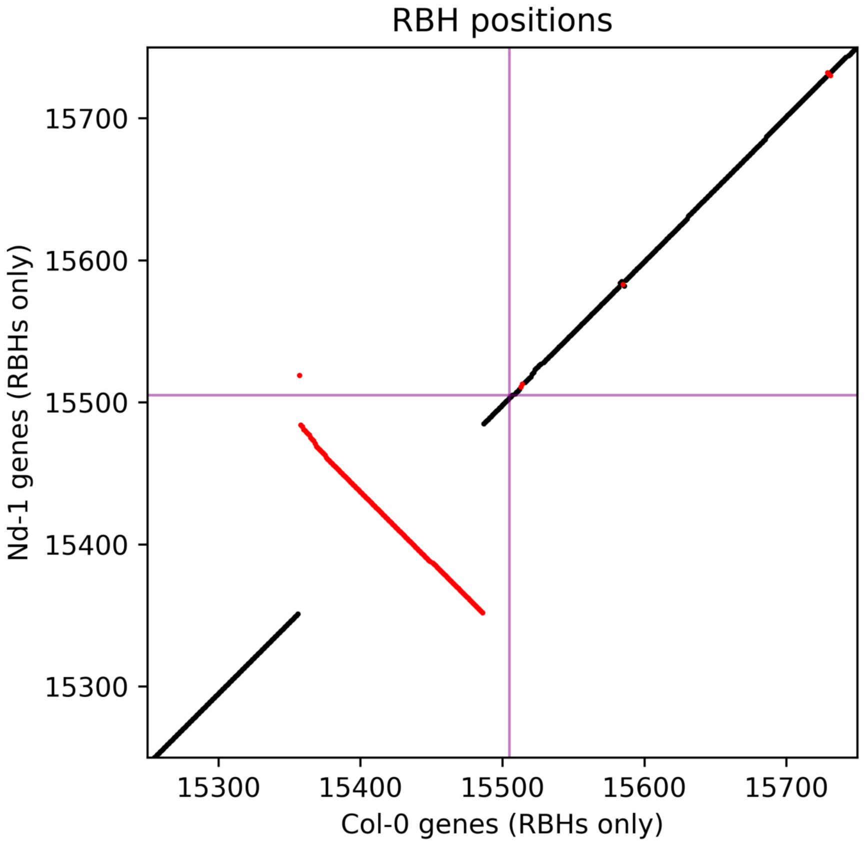

### AdditionalFile27

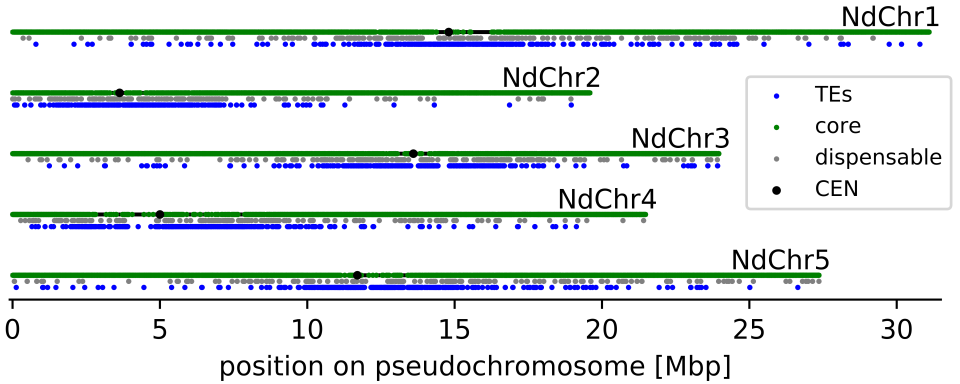

### AdditionalFile28

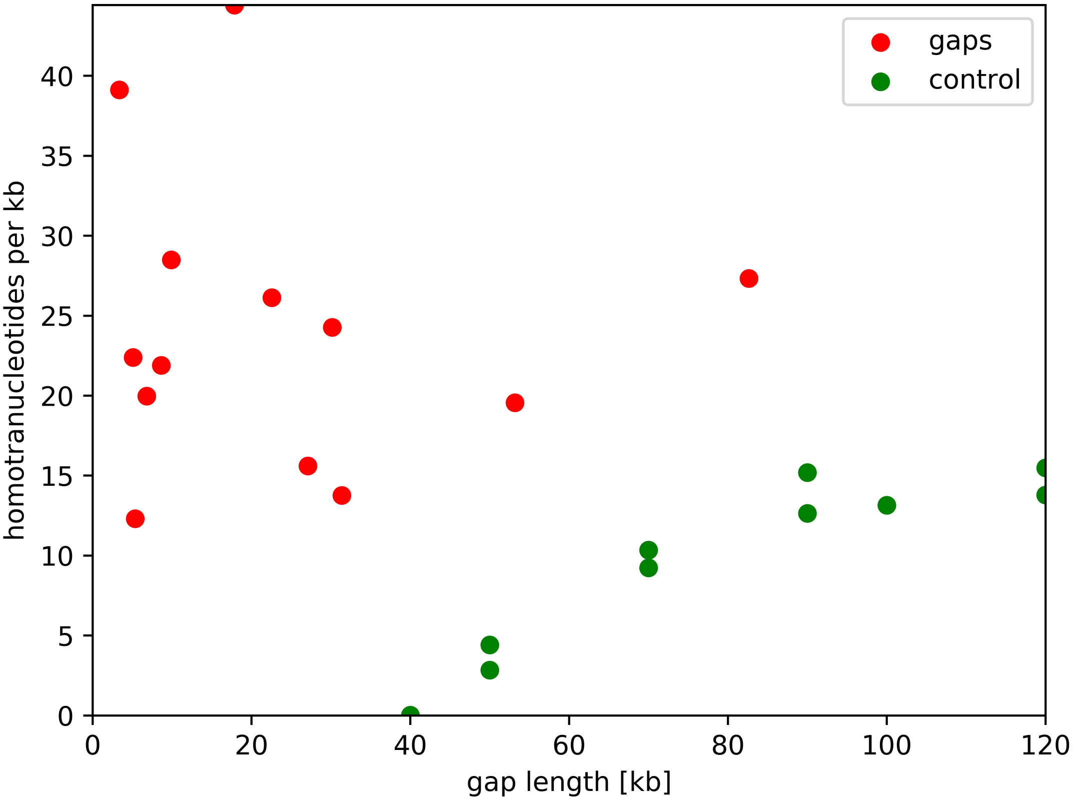
